# Neutrophilic granule protein is a novel murine LPS antagonist

**DOI:** 10.1101/733352

**Authors:** Jaewoo Hong, Peng Qu, Todd R. Wuest, P. Charles Lin

**Affiliations:** Cancer and Inflammation Program, Center for Cancer Research, National Cancer Institute, Frederick, Maryland, 21702, USA

**Keywords:** inflammation, inhibitor, lipopolysaccharide (LPS), neutrophil, protein-protein interaction, small molecule

## Abstract

NGP was previously reported as a granular protein of neutrophils in mouse but the function has not been known clearly. We found the presence of the possible signal peptide in NGP and we hypothesized this protein is being secreted to the blood stream. Since this protein has sequence similarity with an antimicrobial protein cathelicidin, we observed the aspect of inflammation of NGP. Interestingly, NGP interacts with the complex of LPS and LBP and blocked the inflammatory function of NGP. This inhibitory effect of NGP was through the inhibition of LPS-LBP complex to interact with TLR4 molecule. Furthermore, the inhibitory function of NGP could be repeated on the inflammatory effect of LPS in both in vitro and in vivo. With these findings, we report NGP is a novel secretory protein to mask LPS and inhibits its function.

## 1 Introduction

Leukocytes contain several antimicrobial proteins and peptides for the defense of the host organism that undertake through either specific and non-specific pathway. Among the best studied are bactericidal- and permeability-increasing protein (BPI) [1; 2], the defensins [3], azurocidin (CAP37) [4] and LL-37 [5; 6].

LPS initiates signaling triggered by the immunological response to Gram negative bacteria. LPS from several different bacterial species or strains initiates acute inflammatory responses in the host that is typical host reaction to infection and immune cell responses to LPS exposure. LPS can initiate a strong immune response and serves as an early warning signal of bacterial infection. LPS is first released from bacterial membranes and vesicles from them by LPS binding protein (LBP) in host serum. LBP then donates LPS to CD14, which can be found either in soluble form or linked to the cell surface by a GPI anchor. CD14 splits LPS complex into monomeric molecules and donates them to the TLR4–MD-2 complex. Aggregation of the TLR4– MD-2 complex after binding LPS leads to activation of further signaling components, including NF-kB and IRF3, and followed by production of pro-inflammatory cytokines [7; 8; 9; 10; 11].

Neutrophilic granule protein (NGP) was initially identified in immature bone marrow cells and promyelocytes. This 19-kDa myeloid granule protein shows 30% homologous to cathelicidin, which is a member of cystatin superfamily [12]. It has been reported that NGP gene expression is controlled by C/EBP-ε and PU.1 transcription factors, which might be like other myeloid genes [13]. However, the molecular function of NGP protein in vitro and in vivo remains still unclear.

## 2 Article types

Original Research

## 3 Material and Methods

### Animals

The mice were maintained in a pathogen-free facility at the National Cancer Institute (Frederick, MD) in accordance with Animal Care and Use Committee regulations. C57BL/6J mice were purchased from the Jackson Lab and NGP deficient mice on the C57/BL6 background were purchased from Knockout Mouse Project Repository (KOMP, Davis, CA) and backcrossed to C57BL/6J. Sex and age-matched mice were used in all the studies. The LAD ligation was performed as described. For the LPS injection, 0.2 mg/kg of LPS was prepared in saline solution and injected intraperitoneally.

### Cell culture and primary cell isolation

EL-4, Raw 264.7, 293T, THP-1 and HL-60 cell lines were obtained from American Type Culture Collection (ATCC) and maintained according to the instructions. Mouse blood cell lines were isolated from the total blood of mice by MACS (Miltenyi Biotec, Germany) following the description. For the overexpression and shRNA, lentivirus of pHage-NGP and NGP shRNA were prepared in 293T cells following the co-transfection of pMD2g and pAX2. The prepared lentivirus was transduced to cells by polybrene. Mouse neutrophil was isolated from the peritoneal cavity 3 hours after injection of 1 ml thioglycollate medium (Sigma-Aldrich, MO). Isolated neutrophils were incubated at 37C for 6 hours and then treated with 10 ng/ml, 100 ng/ml, 1μg/ml, 10 μg/ml LPS overnight. NGP conditional medium was collected 2 days after transfection of NGP to 293T cells by Fugene HD (Promega). 10 ng/ml of LPS was mixed with conditional medium for 30 minutes and then treated to THP-1 cells.

### qRT-PCR

qRT-PCR was performed using total RNA isolated on RNeasy Quick spin columns (QIAGEN, CA). One μg of total RNA was used to perform reverse transcriptase–polymerase chain reaction (RT-PCR) using iScript supermix (Biorad, Hercules, CA). The sequence of PCR primers used are: NGP, 5’-AGACCTTTGTATTGGTGGTGGC-3’ and 5’-GGTTGTATGCCTCTATGGCTCTA-3’; β-actin, 5’-CTGTCCCTGTATGCCTCTG-3’ and 5’-ATGTCACGCACGATTTCC-3’. Values are expressed as fold increase relative to β-actin and analyzed with CFX manager (Biorad, CA). All primers were purchased from Sigma.

### Generation of NGP antibody

The cDNA sequence of the mature NGP was cloned to pProEx/HT (ThermoFisher) vector. NGP construct was transformed to BL21-Codon plus (Agilent) and induced with 6 mM IPTG for 3 hours. The expressed protein was purified by Talon metal affinity chromatography (BD Biosciences, CA) following the description. The purified recombinant protein was used as the antigen and immunized to rabbits. The immunization and preparation of the antibody was made by Rockland, PA following the company’s protocol.

### Immunoprecipitation and western blot

Conditional medium of NGP expressing THP-1 cells, HL-60 or 293T cells were used for immunoprecipitation by Myc antibody (Cell Signalling, MA). LPS, LBP and NGP conditional medium was mixed at 4C for 30 minutes and then NGP was pulled down with myc antibody. The IP product was propagated to Western blotting for myc tagged NGP detection. The total product of IP was propagated for western blot using Myc antibody. NGP overexpressing THP-1 cells were lysed by 1X sample buffer and then western blotting was performed by myc antibody (Cell Signaling). For NGP western blot, total lysate of mouse cells, mouse tissues, serum or peritoneal neutrophils were used and probed by NGP antibody we generated. IκBα and p-p38, p38 were detected from LPS treated THP-1 cells with or without NGP conditional medium. The antibodies were from Cell Signaling.

### Immunostaining and confocal microscopy

THP-1 cells and HL-60 cells were transduced with NGP-myc lentivirus. Cells were fixed with paraformaldehyde and stained against Golga3, Calreticulin and NGP. Golgia3 and Calreticulin antibodies were obtained from Abcam. Confocal microscopy was performed with LSM-780 (Zeiss, Oberkochen, Germany) and analyzed with ImageJ (NIH, Bethesda, MD).

### Sandwich and direct ELISA

Mouse TNFα, IL-1β and IL-6 were detected by ELISA duo sets (R&D Systems) following the manual. For direct ELISA of LPS receptors, recombinant MD2, CD14 or MD2/TLR4 complex was coated on an ELISA plate overnight at 1μg/ml concentration. Plate was blocked by 5% BSA containing PBS and mixture of 100 ng of biotinylated LPS, 100 ng of recombinant LBP and the conditional medium of NGP or empty vector from 293T cells for 2 hours. Biotinylated LBP was detected by TMB solution (Sigma).

### Molecular cloning

#### Reporter assay

pGluc-NF-κB construct was obtained from Genecopoeia (Rockville, MD). pSV40-CLuc control plasmid was obtained from NEB (Ipswich, MA). These constructs were co-transfected to 293 cells by FugeneHD (Promega). Transfected cells were attached in a 96-well plate and then 0, 1, 10 or 100 ng/ml of LPS was treated overnight. The culture supernatant was projected to perform luciferase assay with Gaussia Luciferase assay kit and Cypridina Luciferase assay kit (NEB) following their description.

### Statistical analysis

All statistical analyses were carried out using Prism 7 (La Jolla, CA). Quantitative variables were analyzed by t-test, one-way ANOVA test. All statistical analysis was two-sided, and p<0.05 was considered statistically significant.

## 4 Results

### Neutrophils express NGP

First, we validated the expression of NGP in the mouse. Total RNA of T cells, B cells, macrophages, monocytes, bone marrow derived neutrophils and blood stream neutrophils were isolated. We compared the mRNA levels of NGP in mouse cells and bone marrow neutrophils and blood neutrophils had high expression of NGP while macrophages and monocytes had lower expression. T cells, B cells and NGP deficient neutrophils had no expression of NGP in mRNA (Figure1A). Next, we also verified the protein expression in mouse neutrophils. We collaborated with Rockland (Limerick, PA) to generate NGP antibody because NGP antibodies are not commercially available. We confirmed the expression of NGP from mouse neutrophils by western blot while EL4 mouse T cell line, Raw 264.7 mouse macrophage cell line and NGP deficient neutrophils had no expression of NGP protein (Figure 1B). The expression of NGP protein was specific in hematopoietic tissue including neutrophils like bone marrow. This protein was not expressed in other tissues such as lung, heart, stomach, ileum, colon, liver, kidney spleen and thymus (figure 1C).

**Figure 1.**
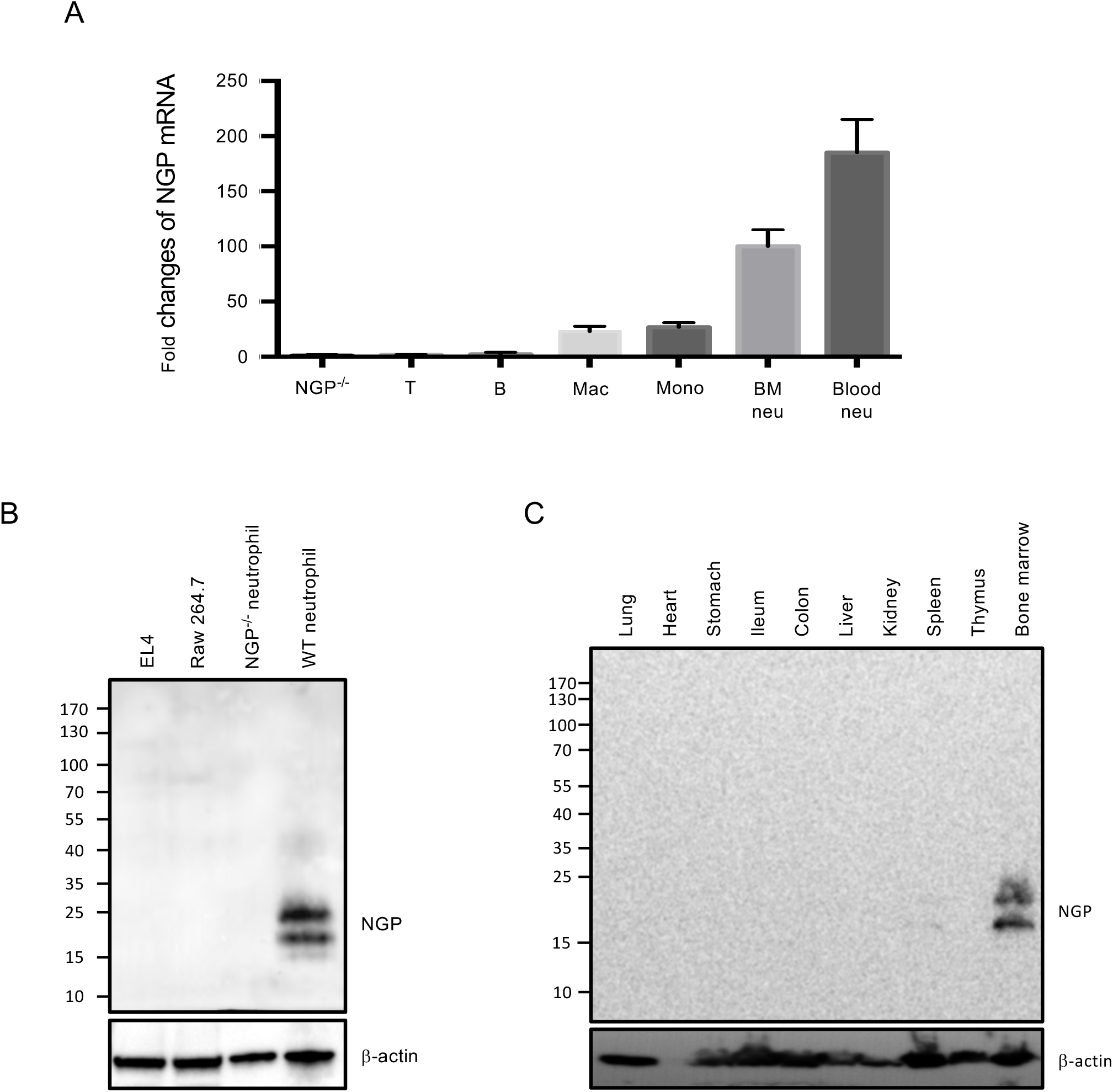
NGP is expressed specifically in mouse neutrophils. A, T cells, B cells, macrophages, monocytes, bone marrow derived neutrophils and circulating neutrophils were isolated from a wild type mouse. mRNA level of NGP was relatively measured by qRT-PCR and compared to NGP deficient neutrophils. B, NGP protein was detected by western blotting from the total lysate of EL4 cells, Raw 264.7 cells, NGP deficient neutrophils and wild type neutrophils by western blot. C, NGP protein was detected by western blotting from the total lysate of mouse tissues. Lung, heart, stomach, ileum, colon, liver, kidney, spleen, thymus and bone marrow were collected from a wild type mouse. Tissue was ground in 1X sample buffer and NGP was detected by western blotting.

### NGP is a soluble secretory molecule

The protein sequence on NGP is anticipated to have a signal peptide with typical 19 hydrophobic amino acids (Figure2A). We hypothesized this protein is simultaneously secreted from cells. NGP was overexpressed in THP-1 and HL-60 cells that have granules and 293T cells to observe the secretion of NGP. The 2-day culture medium of these cells had NGP proteins in the medium without any special stimulation (Figure 2B). Interestingly the location of NGP was ER than granular structures in HL60 cell and THP-1 cells overexpressing NGP (Figure 2C). In addition, when NGP overexpressing THP-1 cells were stimulated by brefeldin A or monensin, NGP protein in the cell was increased. Furthermore, when post-translational modification in ER was blocked by brefeldin A, NGP protein had smaller molecular size than in vehicle or monensin treatment (Figure 2D). The location of endogenous NGP in peritoneal neutrophils was in ER like NGP was overexpressed (Figure 2E). When peritoneal neutrophils were stimulated by brefeldin A or monensin, NGP protein in the cell was increased and the secretion of NGP was reduced (Figure 2D).

**Figure 2.**
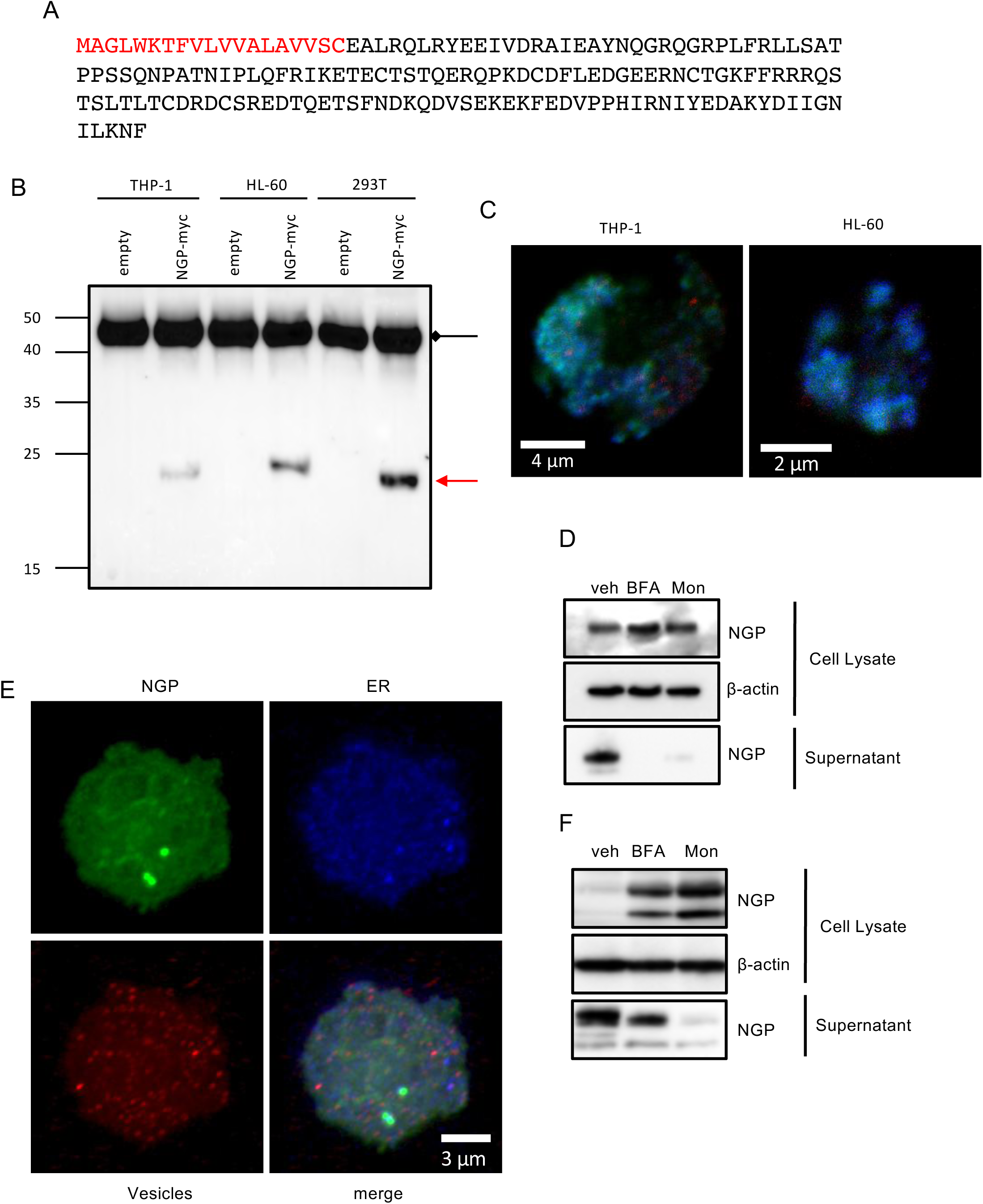
NGP is a soluble secretory molecule. A, The amino acid sequence of NGP. Red letters stand for the area of signal peptide (red). B, myc-tagged NGP was pulled down from NGP overexpressed THP-1, HL-60 or 293T cells. Myc-tag was detected by Western blotting and red arrow shows NGP-myc band. Black arrow shows IgG heavy chain. C, NGP overexpressing HL-60 (upper panel) and THP-1 (lower panel) were stained against Golga3 (red), Calreticulin (green) and NGP (blue). The stained cells were analyzed by confocal microscopy. D, NGP-myc overexpressing THP-1 cells were treated by vehicle, 1X Brefeldin A or 1X Monensin (Biolegend) for 3 hours. Myc-tagged NGP was detected from the total lysate and supernatant by western blotting. E, Mouse peritoneal neutrophils were stained against NGP (green), Calreticulin (blue), lysobrite for granules (red). The stained cells were analyzed by confocal microscopy. D, Mouse peritoneal neutrophils were treated by vehicle, 1X Brefeldin A or 1X Monensin (Biolegend) for 3 hours. NGP was detected from the total lysate and supernatant by western blotting.

### NGP is induced by LPS stimulation

Neutrophils were isolated from mouse peritoneal cavity and 100 ng/ml or 500 ng/ml of LPS was treated for 6 hours to these cells. We detected the expression of NGP protein by western blot and observed NGP protein level is increased when mouse neutrophil is stimulated by LPS (Figure 3A). Next, we isolated mouse serum after stimulation with LPS. NGP protein was detected in the mouse serum 3 hours after mice were stimulated with LPS while NGP deficient mouse did not show any NGP protein in the serum (Figure 3B). The function of secreted NGP in inflammation was assessed from the mouse serum. We injected 0.2 mg/kg of LPS and collected mouse blood to isolate mouse serum. The collected serum was propagated to perform ELISA of TNFα, IL-1β or IL-6. The levels of all TNFα, IL-1β and IL-6 were increased when NGP was knocked out in mice (Figure 3C).

**Figure 3.**
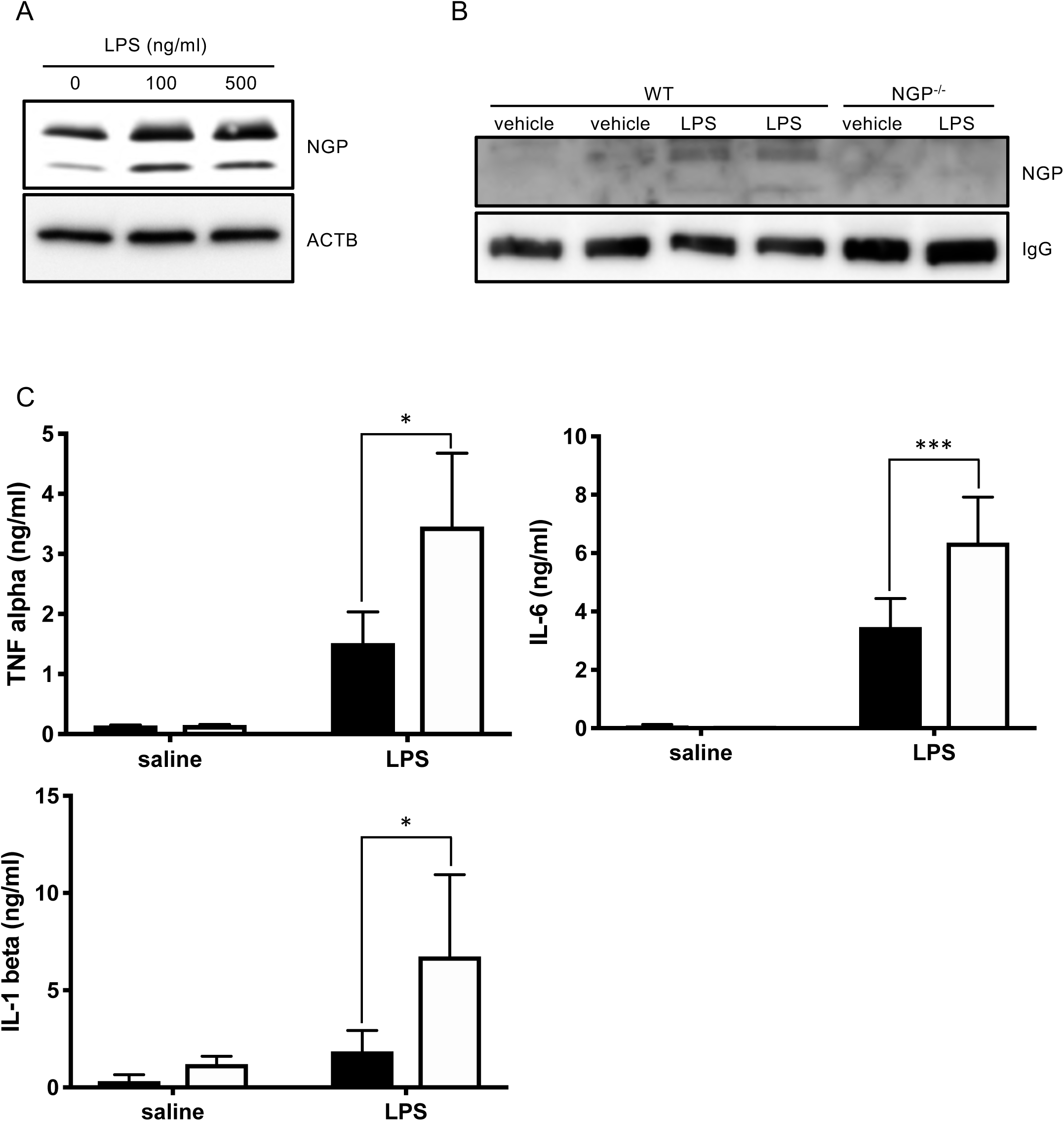
NGP is induced by LPS. A, Mouse peritoneal neutrophils were treated by vehicle, 100 ng/ml or 500 ng/ml of LPS for 3 hours. NGP was detected from the total lysate by western blotting. B, 1 mg/kg of LPS was injected to wild type mouse or NGP deficient mouse intraperitoneally. Serum was isolated from each animal 5 hours after injection. NGP was probed from the serum by western blotting. C, 0.2 mg/ml of LPS was injected to wild type (black bar) or NGP deficient (empty bar) mice. Mouse serum was collected before injection of LPS, 1 hour after injection for detection of IL-1β and IL-6 and 3 hours after injection to detect TNFα. TNFα, IL-1β and IL-6 were detected by ELISA.

### NGP interacts with LPS binding protein (LBP)

The next question was how NGP inhibits the LPS function. We mixed LPS and/or LPS binding protein (LBP) in the conditional medium of empty vector or myc-tagged NGP from 293T cells. Then we pulled down NGP with the tag antibody. NGP was pulled down from the overexpressed conditional medium and interestingly, LBP protein was pulled down with NGP only in presence of LPS. This shows NGP pulls down LBP only when LPS exists (Figure 4A). We prepared wild type and 5 different NGP mutant clones with C-terminal myc-tag to identify the binding site of LPS-LBP complex and NGP (Figure 4B). The culture medium of wild type or NGP mutants were mixed with LBP and LPS and then NGP was pulled down with myc tag. Wild type NGP and C-terminal 20, 60 and 90 amino acid deletion mutants pulled down LBP (Figure 4C); however, the deletion mutant of Glu_20_-Leu_47_ did not pull down LBP unlike wild type NGP or other mutants (Figure 4D). Next, we screened which of LPS receptors is being blocked for LPS-LBP complex to bind. We coated recombinant MD2, CD14 TLR4 or TLR4/MD2 complex in the ELISA plate and added the mixture of LPS and NGP in control or NGP conditional medium. LPS binding to the coated receptor component was inhibited with NGP conditional medium when TLP4 or TLR4/MD2 complex were coated unlike when MD2 or CD14 was coated, which shows NGP inhibits the binding of LPS-LBP complex to TLR4 (Figure 4E). Since the binding of LPS-LBP with CD14-MD2-TLR4 complex is inhibited by soluble NGP molecule in extracellurar matrix, the downstream signaling of cannot be made successfully from the first (Figure 5).

**Figure 4.**
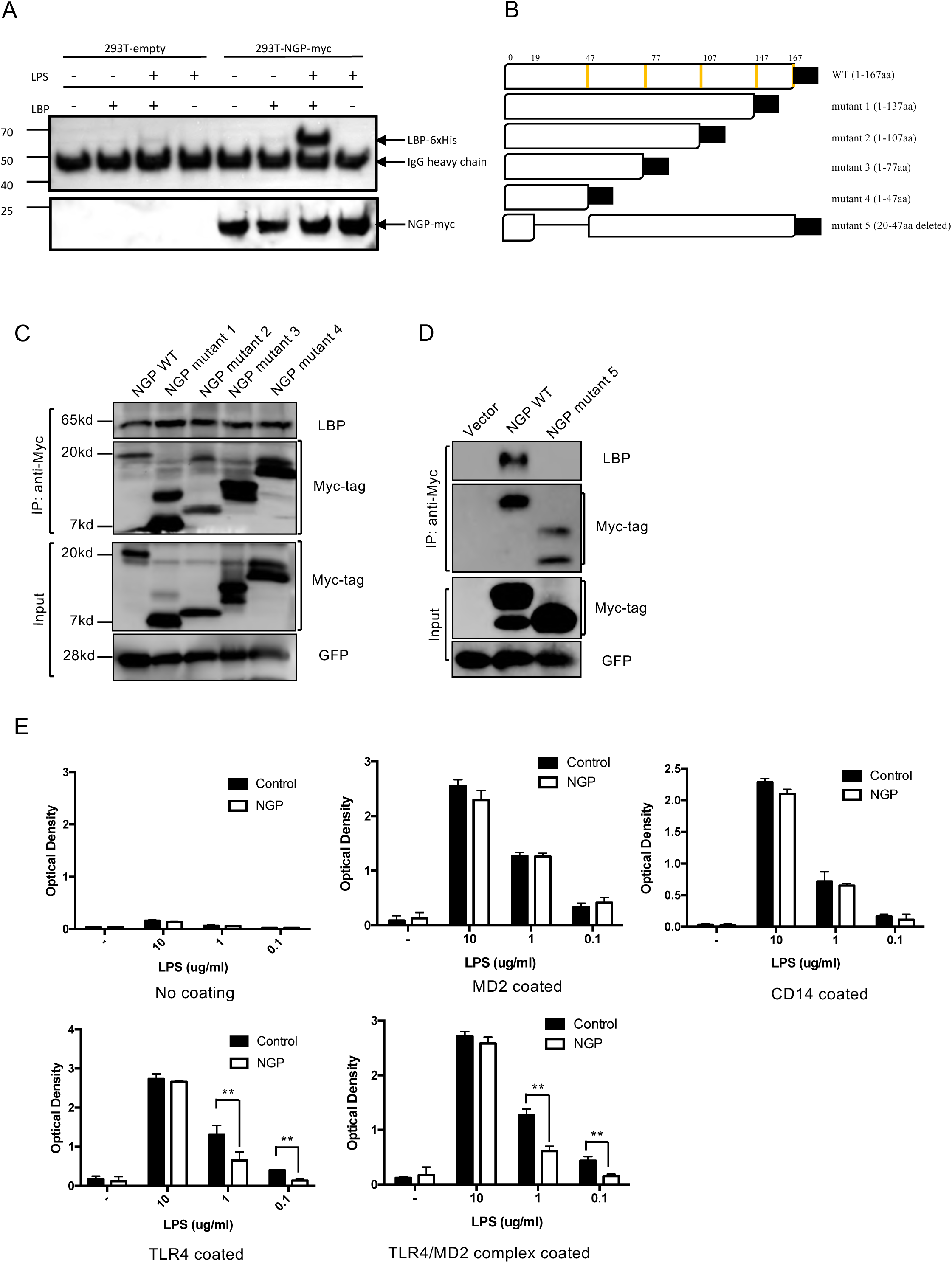
NGP masks LPS and inhibits LPS receptor binding. A, Conditional medium of NGP or empty vector was mixed with 100 ng of recombinant LBP-His and/or 100 ng of LPS for 30 minutes on ice. LBP or NGP was detected by western blotting. B, Five different NGP mutant constructs were prepared to identify the binding site of LPS-LBP complex on NGP. Amino acids are deleted from the C-terminus in mutant 1-4 as designated. Mutant 5 has deletion of 28 amino acids after signal peptide. C and D, Wild type NGP or deletion mutants of NGP were mixed with 100 ng of LPS and 100 ng of LBP for 30 minutes on ice. NGP was pulled down with myc-tagged NGP and LBP and myc-tagged NGP were detected by western blotting from both IP product and input. E, Recombinant MD2, CD14, TLR4 or TLR4/MD2 complex was coated on an ELISA plate. Biotinylated LPS was mixed with 100 ng of LBP in designated concentration for 30 minutes on ice. The mixture of LPS and LBP was added to the receptor coated plate for 2 hours. Bound LPS was visualized by TMB solution and analyzed by ELISA reader.

**Figure 5.**
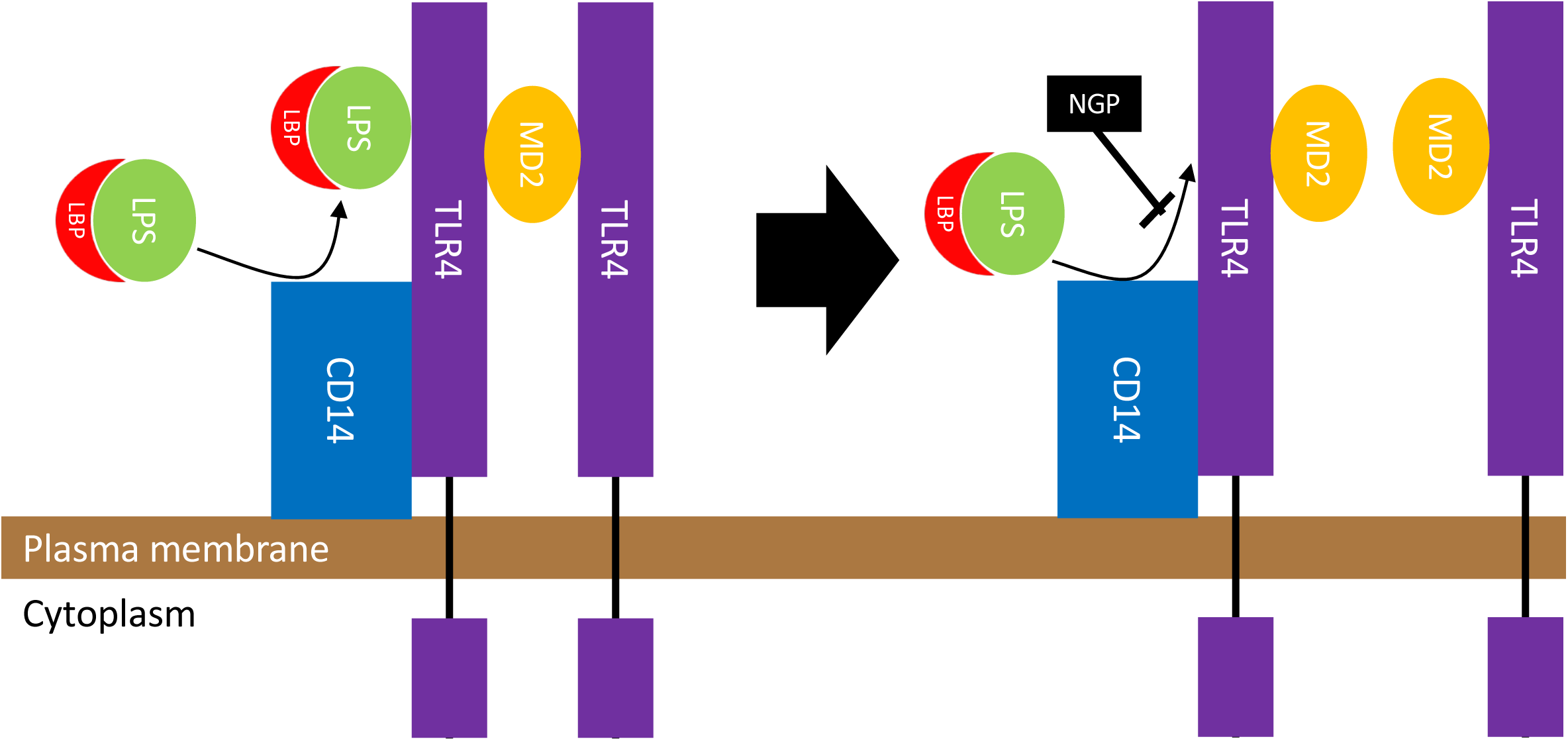
NGP masks LPS-LBP complex. The complex of LPS and LBP stimulates the LPS receptor complex of CD14, TLR4 and MD2. NGP binds to the LPS-LBP complex and masks the complex to interact with its receptor. LPS-LBP complex fails to bind to TLR4 from CD14 and followed by the null downstream signal of TLR4.

### NGP inhibits LPS activity

We transfected NGP or empty vector in 293T cells to collect the conditional medium of NGP 2 days post-transfection. We mixed conditional medium of empty vector and NGP vector in several ratio and then added this medium to THP-1 cells. Then we treated 10 ng/ml of LPS to THP-1 cells and incubated overnight. We harvested the LPS treated medium and measured IL-8 secretion by ELISA. IL-8 was highly secreted in the empty vector conditional medium but it was remarkably decreased when NGP conditional medium was mixed in high concentration. This suppression of IL-8 was lower as the concentration of NGP conditional medium was low (Figure 6A). We further assessed whether the LPS mediated downstream signals are affected by NGP treatment. The conditional medium of empty vector or NGP vector was mixed with vehicle or LPS and then treated to THP-1 cells. As the result, LPS induced the degradation of IκBα in the cell within empty vector medium while NGP medium kept the level of IκBα high and decreased the phosphorylation of NF-κB p65 (Figure 6B). In similary way, the phosphorylation of p38 MAPK was increased high when LPS was treated in empty vector medium but the phosphorylation was lowered when cells were in NGP conditional medium (Figure 6C). The conditional medium of NGP inhibited the activity of LPS mediated induction of NF-κB. LPS was mixed in different concentrations and then treated to 293 cells transfected with Gaussia luciferase NF-κB construct. The luminescence of cells with NGP medium showed inhibited activity of the reporter (Figure 6D). Next, we collected mouse peritoneal neutrophils from wild type or NGP deficient mice. We treated LPS to these neutrophils in different concentrations overnight and then collected culture supernatant. The level of inflammatory cytokine, IL-6 was measured by ELISA. The level of secreted IL-6 increased as the concentration of LPS was higher and IL-6 was secreted much more when the neutrophils were collected from NGP deficient mice (Suppl Fig 1C).

**Figure 6.**
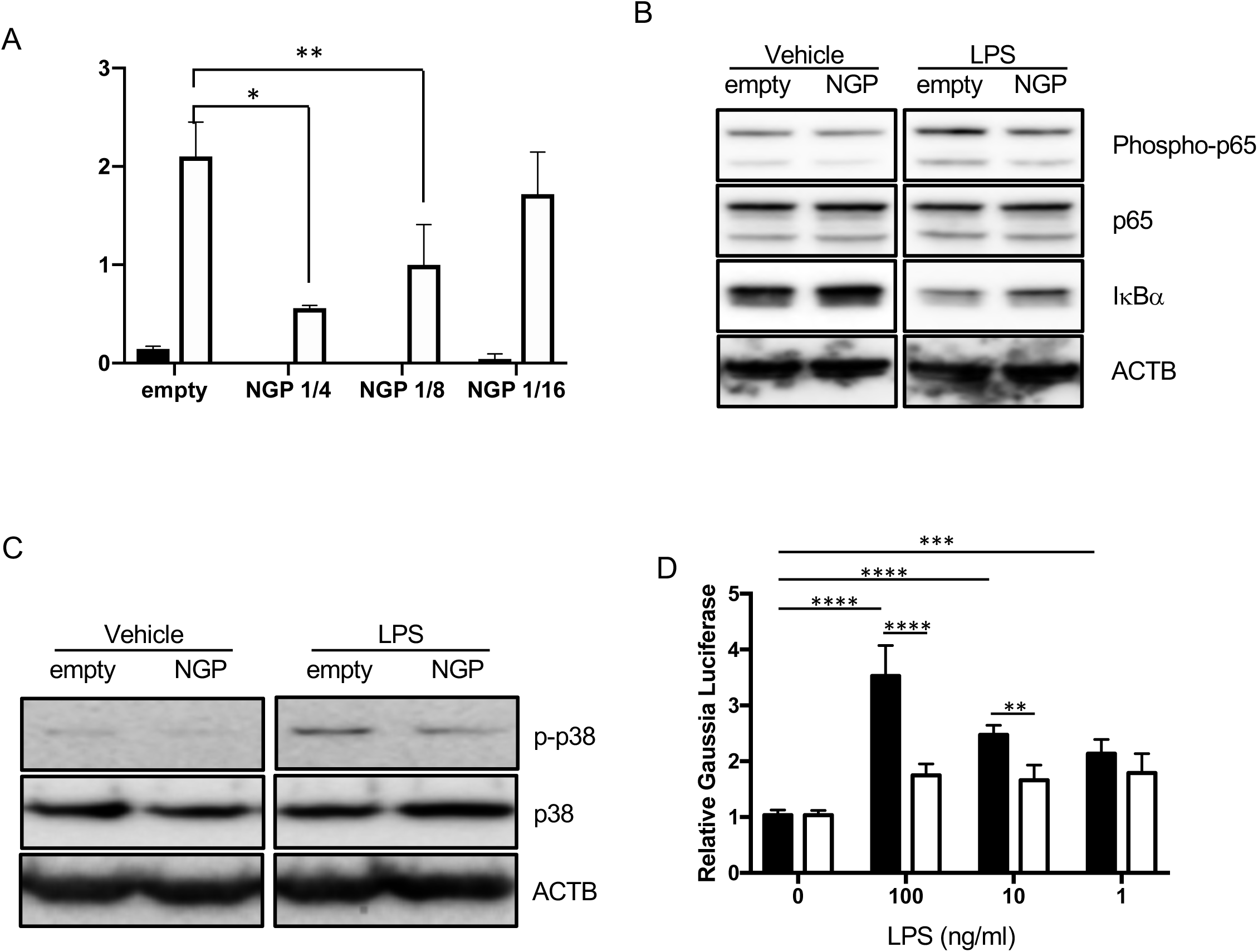
NGP blocks the activity of LPS. A, THP-1 cells were treated by vehicle (black bars) or 100 ng/ml of LPS (blank bars) for 16 hours in the mixture of NGP conditional medium and empty vector conditional medium in ratio of 0, 1/4, 1/8 and 1/16. IL-8 was detected from the culture medium. B, THP-1 cells were treated by vehicle or 10 ng/ml of LPS in empty vector or NGP conditional medium for 30 minutes. Phospho-NF-κB p65, NF-κB p65 and IκBα were detected by western blotting. C, THP-1 cells were treated by vehicle or 10 ng/ml of LPS in empty vector or NGP conditional medium for 30 minutes. Phospho-p38 MAPK and p38 MAPK were detected by western blotting. D, The conditional medium of empty vector (black) or NGP (blank) was treated overnight. LPS was mixed in different concentrations and then treated to 293 cells transfected with Gaussia luciferase NF-κB construct. The luminescence of cells with NGP medium showed inhibited activity of the reporter.

## 5 Discussion

NGP was first discovered in mouse neutrophils and the function has not been known. The closest orthologue in human is cathelicidin, which is an antimicrobial protein and we hypothesized this protein has a role in bacterial infection. We first validated this molecule is expressed in mouse neutrophils. Several lineages of mouse blood cells were isolated and assessed the expression of NGP mRNA in those cells. As it has been known, the level of NGP was solely high in bone marrow or circulating neutrophils and the level was higher in circulating neutrophils than bone marrow neutrophils.

We had a limitation to study NGP molecule because the antibody against NGP is not commercially available. The antibody against NGP had to be generated so we expressed and purified recombinant NGP protein from *E.coli*. This recombinant protein was used as the antigen to generate NGP antibody from a rabbit in collaboration with Rockland. Through the validation of the antibody by antigen-direct ELISA and western blot using the antigen, the antibody showed high specificity and affinity to both recombinant and endogenous NGP protein. We could detect NGP protein in mouse neutrophils finally using this NGP antibody. As we expected, NGP protein could be easily detected from mouse neutrophils unlike other mouse leukocyte cell lines or NGP deficient mouse neutrophils. Furthermore, NGP expression could be detected from bone marrow, which has high portion of PMN cells than other mouse organs like lung, heart, stomach, intestine, liver, kidney, spleen and thymus. This confirmed NGP protein is neutrophil specific protein. Interestingly, NGP protein had double band under western blot, which may be the result of post-translational protein that is vigorously modified and secreted.

The amino acid sequence of NGP protein has typical 19 hydrophobic amino acids on its N-terminal, so we hypothesized this protein is being secreted out of cells. We validated the secreted NGP protein in the culture medium when NGP is overexpressed from THP-1 cells and HL-60 cells that have granules and in 293T cells. Next, we overexpressed NGP in THP-1 cells and blocked the protein maturation in ER or golgi with Brefeldin A or Monensin consequently. When these steps were blocked, the protein level of NGP was kept high in the cell, not being secreted. Furthermore, the molecular size of NGP was lower when ER process was blocked, which showed this protein is being modified in ER, which also is a classic evidence this protein goes through secretory protein processing.

We also wondered if this protein is being secreted into the blood stream and whether this protein is being affected by bacterial stimulation such as LPS treatment in vivo. We first observed the protein level of NGP was increased when primary mouse neutrophil was stimulated by LPS in dose dependent manner. Next, the protein in the circulating mouse blood could be detected in isolated serum of mouse when mice got injection of LPS. NGP is secreted into the blood stream in response to LPS, then we wondered whether NGP affects any function of LPS. Interestingly, the increase of proinflammatory cytokines such as TNFα, IL-1β and IL-6 in NGP deficient mice by the stimulation with LPS was much higher than wild type mice. This showed NGP is secreted into blood stream by LPS stimulation and lowers the inflammatory effect of LPS.

The inhibition of LPS mediated proinflammatory cytokine activation was also inhibited when the conditional medium of NGP was mixed with LPS in vitro. Logically the downstream signaling like p38 MAPK and NF-κB by LPS stimulation on CD14-TLR4-MD2 pathway was downregulated when NGP was upregulated in the cell. Furthermore, the isolated neutrophils of NGP deficient mice showed much more sensitive response to LPS unlike wild type mice. This shows NGP is an inhibitor of LPS not only in in vivo but also in vitro.

The binding of NGP to LPS-LBP complex is the key mechanism to suppress LPS activity. NGP binds to LBP in presence of LPS. NGP did not bind to LBP when LPS was not mixed with LPS. We wondered which site of NGP binds to LPS-LBP complex, so we prepared several different deletion mutants of NGP protein. This revealed amino acids between Glu_20_-Leu_47_ of NGP bind to LBP-LPS complex. In addition, the interaction of LPS-LBP complex with LPS receptor could be inhibited by NGP. A component of LPS receptor, TLR4 binds to LPS-LBP complex like other components, MD2 and CD14 in dose dependent manner but when LPS-LBP complex is mixed with NGP conditional medium, the interaction of TLR4 with LPS-LBP complex was significantly inhibited. This shows NGP is an antagonist of LPS-LBP complex and inhibits the interaction of this complex to TLR4.

In this study, we found the signal peptide sequence of NGP molecule that secretes this protein from cells. NGP captures LPS-LBP complex and inhibits the interaction with TLR4 which blocks the downstream activation of TLR4 pathway. NGP is not the only inhibitor of LPS but this molecule blocks the function of LPS at the highest level of LPS signaling. In history, the fight of host immune system against bacterial infection has been always challenging and will be continued forever. NGP may not enough to overcome endotoxemia or endotoxin shock of the host; however, NGP gives us a direction to cure an additive or substitutive therapy of bacterial infection.

## 7 Author Contributions

JH, TRW and PCL designed experiments; JH performed experiments; JH and TRW analyzed data; JH and PCL prepared the manuscript

## 9 Acknowledgments

This is a short text to acknowledge the contributions of specific colleagues, institutions, or agencies that aided the efforts of the authors.

