## Supplementary figures and images for "Neutrophilic granule protein is a novel murine LPS antagonist"

### suppl fig

Supplementary Figure 1

A

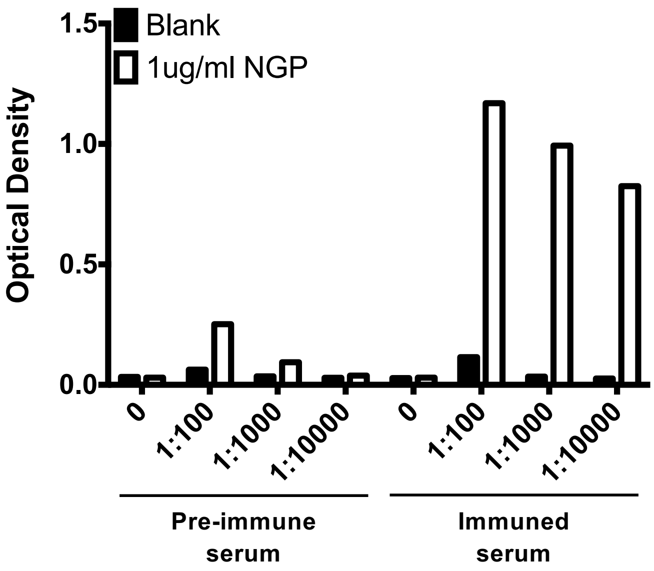


B

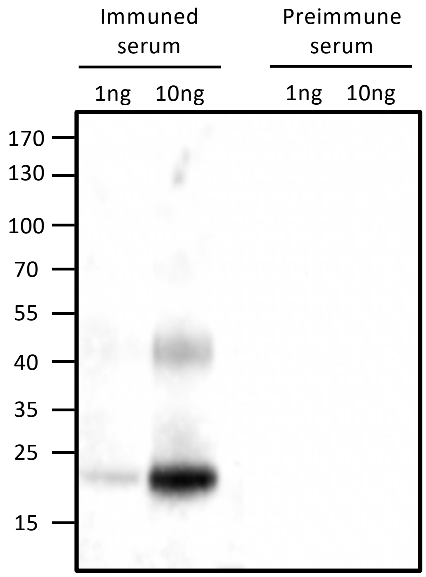

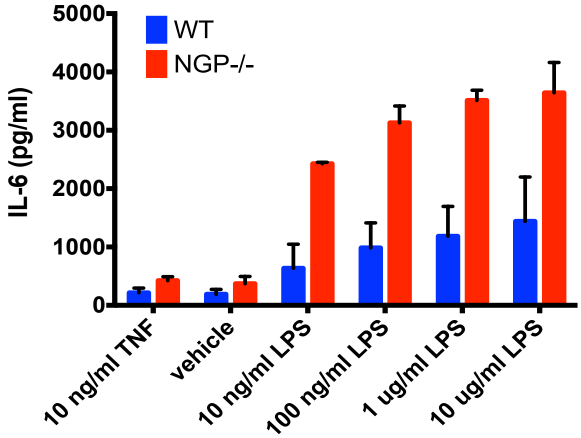
