## Supplementary material for "Neutrophilic granule protein is a novel murine LPS antagonist": suppl legend

**Supplementary Figure 1.** A, 1 μg/ml of recombinant NGP was coated on an ELISA plate. Preimmune serum and immuned serum of NGP from the rabbit was added in designated concentration for 2 hours at room temperature. The bound antibody was measured by donkey anti-rabbit IgG-HRP antibody and visualized by TMB solution. The signal was analyzed by ELISA reader. B, 1 ng or 10 ng or recombinant NGP was detected by western blotting with NGP immuned serum or preimmune serum.

**Supplementary Figure 2.** Peritoneal neutrophils of a wild type mouse or NGP deficient mouse were isolated and stabilized in 10% FBS containing RPMI for 6 hours. LPS of designated concentrations was treated overnight to neutrophils. IL-6 was detected from the culture medium by ELISA.
